# Steering Before Selection: Energy-Limited Developmental Control as a Phenotypic Capacitor, a Pre-Specified Simulation Study

**DOI:** 10.64898/2026.11.03.763948

**Authors:** Mohammed Qasim, Faisal Fayyaz Qureshi

## Abstract

Living systems meet environmental change at several speeds: fast physiological control during development, slower learned memory, and genetic change through selection. We asked whether developmental control, which we interpret as bioelectric and which carries a continuous energy cost, steers evolution, and we report what survived twenty-two versions of individual-based simulation, pre-specified from version 11 onwards. After a permanent environmental change, sensing absorbed the change within about 25 generations and genes took over through selection alone 340 to 450 generations later in every seed, a result replicated in a fresh pre-specified test on 20 new seeds. Control did not guide genetic change; it concealed genetic variation from selection. Control hid three to six times more harmful load than no control, and an energy shortfall released it; brief release outperformed prolonged release, and release helped only when control re-locked around the new form. The net value of release depended mainly on whether the concealed harmful load was kept in check: without reversal of harmful mutations it was a cost at 500 individuals (mean fitness 0.103 against 0.122 without control), whereas with reversal it was a benefit at every population size from 250 to 4,000 (0.632 against 0.489 at 2,000). A test that separated the factors showed that population size alone did not reverse the result and that the cost lay in the concealment itself, with release recovering part of it. We conclude that energy-funded developmental control buys time while selection rewrites the genes, conceals rather than directs variation, and acts as an energy-switched phenotypic capacitor whose value depends mainly on whether the harmful load it conceals is kept in check. The models are abstract and contain no voltages or channels, so the bioelectric reading is an interpretation and all results are model results. We set out predictions for testing these claims in developing embryos.

## 1. Introduction

Natural selection is blind, yet organisms are not passive while it acts. During development, tissues correct towards a target form using patterns of membrane voltage that can be measured in living embryos (Adams and Levin, 2012), and simulations suggest that this developmental competency changes how evolution proceeds by masking the raw fitness of genomes (Shreesha and Levin, 2023). This work began from a simple question: does such control merely buffer development, or does it steer evolution, and at what energetic price?

We framed the question around three roles that bioelectric control might play during development. Energy flow is the **fuel** that pays for maintaining voltage. A ratchet that keeps what works acts as **memory**. Self-correction towards a target form is the **steering**, which acts on the body before natural selection acts on the genes. We called the idea *steering before selection*.

Three bodies of theory bear directly on it. Work on phenotypic plasticity shows that the timescale of environmental change and the reliability of cues decide whether organisms rely on fixed genes, plastic responses or a compromise between them (Botero et al., 2015), that plastic responses can come first and be genetically assimilated later when plasticity is costly (Scheiner et al., 2017), and that organisms can integrate genetic, parental and environmental information (McNamara et al., 2016). Work on cryptic genetic variation shows that variation concealed from selection can be released by stress or by loss of a buffer such as the chaperone Hsp90 (Rutherford and Lindquist, 1998; Paaby and Rockman, 2014), as in cavefish, where a shift to cave water appears to have released hidden variation in eye size (Rohner et al., 2013). Finally, whether such release benefits populations is contested, with theory indicating that it can be favoured only above a minimum population size (Masel, 2005, 2006).

In this paper, “bioelectric control” is the interpretation we give to the control parameter of the simulations; the models themselves contain no voltages or ion channels (sections 3 and 6.5). The paper concerns how control and selection interact, not why life persists or whether it is more than material; those questions lie outside the models and outside the claims. It reports a programme of twenty-two simulation versions that tested the original idea against these alternatives, together with the hypothesis that survived. The original claim that control steers genetic change did not survive; what did survive is a more specific account of what energy-funded control does and when it pays.

## 2. Hypothesis

The hypothesis supported by the full programme is as follows.

During development, energy-funded bioelectric control holds the body to a working form, absorbing environmental change faster than genes can, while selection gradually writes lasting changes into the genes. This control does not guide genetic change; it conceals genetic variation, harmful and useful alike, and it is maintained only where it pays, being cut back where there is nothing to correct. When energy runs short, control weakens automatically and releases the concealed variation; a brief release speeds adaptation, whereas a prolonged one is costly. Whether release is a net benefit or a net cost depends mainly on whether the concealed harmful load is kept in check (in the model, by reversal of harmful mutations) and, more weakly, on population size; where there is a net cost, it lies in the concealment itself, and release recovers part of it.

(*Bioelectric* is the interpretation of the model’s control parameter; the simulations do not model voltage. See sections 3.2 and 6.5.) The hypothesis applies to the developmental window, while the body is being built, and not to finished adults.

The hypothesis changed substantially under testing, and the changes are part of the result.

| **Original claim** | **Outcome** | **Versions** |
|---|---|---|
| Steering guides genetic change | Not supported: steering concealed genetic variation from selection | 1, 14, 15 |
| Memory forms a middle rung between sensing and genes | Not supported unless memory is passed between generations, as by culture | 14–16 |
| Each layer takes the largest energy share where it matters | Not supported | 12 |
| Fast control absorbs change; genes follow through selection | Supported | 15, 16 |
| Control is cut where it is not needed and pays only under damage or change | Supported | 4, 5, 11–13 |
| Concealed variation is released when energy runs short | Supported; net value depends mainly on whether harmful load is kept in check, and more weakly on population size | 17–21 |
| Net value of release depends on population size | Not supported as the main factor at sizes 250 to 4,000 | 19–21 |

## 3. Methods

The programme used two individual-based models: a layered model of developmental control for Part 1 (versions 1 to 16, reported here through its final form, versions 15 and 16, plus the re-test of version 16 as version 22) and a population-genetic model of concealment and release for Part 2 (versions 17 to 21).

### 3.1 Part 1: the layered model

Each of 40 individuals is a 10 × 10 grid of cells whose values make up its body form. The environment demands one of two target forms, a smooth gradient (A) or a checkerboard (B). Each individual carries an inherited genetic template, which changes only through mutation and selection, and a learned memory, which is reset at birth except for a small carried-over fraction (5% by default). Five heritable dials, each between 0 and 1 and free to evolve, set how much the individual relies on its genes, its memory and its senses, how strongly its steering engine corrects the body, and (in one experiment) how far behaviour reshapes the environment.

A generation is one lifetime of ten episodes. At birth the body is built from the genetic template with developmental noise, and random patches of damage are applied. In each episode the individual receives a noisy cue of the current target (cue noise sd 0.3); if it has invested in sensing, it updates its memory towards the cue. The steering engine then moves the body, over five steps, towards a weighted combination of the genetic template, the memory and the cue, the weights being set by the dials. Following the restructuring proposed during the programme, steering is the engine driven by the other three sources rather than an independent competing layer.

Every capacity costs energy: an upkeep cost for each dial, a holding cost proportional to steering strength, a work cost proportional to the correction actually performed, and a cost for writing memory. Genes carry their own cost of accuracy, since a higher gene dial lowers the mutation rate of the template. Fitness in each episode is the negative squared mismatch between body and target, minus the energy spent, and the top third of the population by lifetime fitness reproduces. Environments either alternate between A and B with a set persistence (1, 10, 100 or 1,000 episodes) or switch permanently from A to B at generation 100. Version 16 added a contribution measure, the share of the correct form supplied by genes, memory and sensing.

### 3.2 Part 2: the capacitor model

Part 2 used a Wright–Fisher model with sexual reproduction and free recombination. Each individual carries 20 additive shape loci, whose sum gives the genetic value G, and (from version 18) 20 defect loci, whose sum gives the harmful load D. Development adds noise to give a raw form R = G + ε, and bioelectric control pulls the expressed form P towards a stored target T:

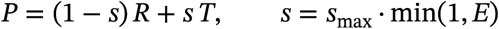

where E is available energy (1 normally, 0.2 under shortfall) and *s*_max_ = 0.8. Nothing in the model decides to loosen control; a shortfall lowers s directly. Control conceals defects in the same proportion, and fitness combines a Gaussian term around the environmental optimum O with a defect term:

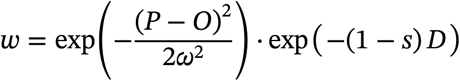

with ω = 1. The energy cost of control multiplies absolute fitness by (1 − 0.1s); it falls equally on all individuals and is reported separately. When energy returns, the target re-locks onto the population’s current form.

| Version | What it added | Population | Mutation inputs | Environment |
|---|---|---|---|---|
| 17 | Concealment and release with an energy switch; neutral shape mutations | 500 | Shape mutation 0.01 per locus, sd 0.1 | One shift of the optimum from 0 to 2 |
| 18 | 80% harmful mutations, shortfall duration, re-locking, random timing | 500 | Harmful effects exponential, mean 0.05 | One shift; shortfalls of 5, 20, 50 or 300 generations |
| 19 | Published mutation figures, control that slowly tracks the current form, repeated changes | 500 | U = 1.2 (Haag-Liautard et al., 2007); gamma shape 0.23 (Eyre-Walker, Woolfit and Phelps, 2006); mutational heritability 5 × 10<sup>−4</sup> (Houle et al., 1996); effects and mutational heritability rescaled by 20, U unchanged | Ten shifts of ±1, one every 150 generations |
| 20 | Larger population, back mutation, a two-trait valley | 2,000 | As version 19 but rescaled by 5; reversion at 10% of the forward rate | As version 19, plus a valley experiment |

Two later versions were run on 20 October 2026 and are reported in sections 4.5 and 5.5. Version 21 reuses the version 20 model unchanged and crosses population size (250 to 4,000), the rescaling factor (Q = 20 or 5) and back mutation (none or 10% of the forward rate), with 20 seeds per cell. Version 22 reruns the version 16 model unchanged on new seeds.

Rescaling multiplied effect sizes and mutational variance by the ratio of a human-like effective population size (about 10,000) to the simulated population, so that selection relative to drift matched the source species.

### 3.3 Procedure

From version 11 onwards every version was pre-specified before its code was run (the written plans are called pre-registrations in the repository file names, but they were not lodged with an external registry): predictions, scoring thresholds, failure conditions and robustness settings were fixed in writing, and every later change was logged with its reason. Each version used ten or twenty seeds per group, a null comparison (for example, a population without control, or with every dial switched off), and robustness checks varying costs, starting values and key parameters. From version 19 onwards a pre-specified settle rule mapped each possible outcome to a verdict, accepted only if it recurred in at least 75–85% of robustness settings. Design-validity checks before scoring were permitted only on the comparison group, so that no fix could be tuned towards a result. Results obtained only in exploratory voice conversations with the AI assistant and never reproduced in text are excluded.

### 3.4 Summary statistics

Groups compared within a seed share that seed, so comparisons are paired. For each comparison we report the number of seeds in which one group exceeds the other and the median of the per-seed differences with its interquartile range (IQR, inclusive quartiles). We add an exact two-sided sign test on the seed counts (null probability 0.5), computed from the saved per-seed values for versions 20 and 21 and from seed counts alone for versions 17 to 19, whose per-seed values were not saved. These p-values are descriptive. The verdicts were fixed in advance by seed-count thresholds; seeds are simulation replicates, so a small p-value shows that a difference is consistent across replicates and not that it would be found in nature; no correction is made for multiple comparisons; and with 20 seeds the smallest attainable p-value is 1.9×10^−6^. Supplementary Table 1 gives the statistics for every comparison.

## 4. Results, Part 1: layered control and the handover to genes (versions 1–16 and 22)

Fast control absorbed environmental change first, and genes took over lasting changes through selection alone, hundreds of generations later. Control did not guide that genetic change.

### 4.1 Control is kept only where it pays (versions 4, 5, 11–13)

Across four versions, sensing was switched off by selection wherever there was nothing reliable to sense, and in a calm, stable world all forms of control were cut back, which parallels the loss of eyes in cave-dwelling animals. Steering paid only where there was damage or environmental change, and under a pump-based energy cost there was an intermediate optimum rather than a simple rule that more control is better. The prediction that each layer would take the largest share of the energy budget in the environment that favours it failed (version 12).

### 4.2 Steering conceals rather than guides (versions 1, 14, 15)

In every version that tested it, steering reduced the visibility of genetic differences to selection rather than directing genetic change. Learned memory never became a leading source of the correct form when it was reset at birth; it contributed 4–7% of the form, rising to 14–20% only when 20% of it was carried over between generations, as culture would allow (versions 14–16). With sensing, memory and steering present, genes took a median of about 312 generations to halve their mismatch after a permanent change, against about 276 generations with genes alone (9 of 10 seed pairs): flexibility slowed genetic adaptation slightly but did not stop it (version 15; reported from the original session and not rerun). A behaviour route, in which behaviour reshapes the environment and genes then follow, succeeded in only 5 of 10 runs and depended on a chosen bonus value (also reported from the original session and not rerun).

### 4.3 Who supplies the correct form depends on how long a change lasts (version 16)

When the environment flickered every episode, genes carried most of the form by settling on a compromise between the two targets, a bet-hedging outcome that was not predicted and depends partly on the baseline used to measure contributions. For changes lasting 10 to 1,000 episodes, sensing led.

| **Persistence of environment (episodes)** | **Genes** | **Memory** | **Sensing** | **Leading source (seeds)** |
|---|---|---|---|---|
| 1 | 0.75 | 0.00 | 0.25 | Genes (10 of 10) |
| 10 | 0.41 | 0.06 | 0.53 | Sensing (8 of 10) |
| 100 | 0.31 | 0.06 | 0.62 | Sensing (10 of 10) |
| 1,000 | 0.44 | 0.07 | 0.49 | Sensing (9 of 10) |

### 4.4 The handover after a permanent change (version 16)

After a permanent switch of the target at generation 100, sensing supplied about 95% of the correct form within about 25 generations. Genes then caught up through selection alone and overtook sensing between generations 438 and 549 in all 10 seeds (median 489, IQR 462 to 510; sign test p = 0.002), that is, 338 to 449 generations after the change; by generation 600 they supplied 65% of the form (memory 6%, sensing 29%). This timescale is of the same order as human lactase persistence, which began to spread roughly 7,500 years ago (Itan et al., 2009), and high-altitude adaptation in Tibetans, with selection estimated to have begun about 9,000 years ago (Zhang et al., 2021); at 25 to 30 years per generation these are of the order of 250 to 300 and 300 to 360 generations, though the published intervals are wide. The correspondence reflects the parameters chosen and is not a validation of the model. Robustness depended on costs and inheritance of memory: genes led in the late window with halved costs (8 of 10 seeds) or 20% carry-over of memory (10 of 10), and sensing led with doubled costs or no carry-over (8 of 10 each). Because the crossover fell inside the pre-specified scoring window, the pre-specified comparison was scored as a tie and the crossover was identified afterwards; it therefore required a fresh pre-specified test, reported in section 4.5. All version 16 figures above were reproduced exactly from the recovered code on 18 October 2026. The trajectories are shown in Figure 1.

**Figure 1.**
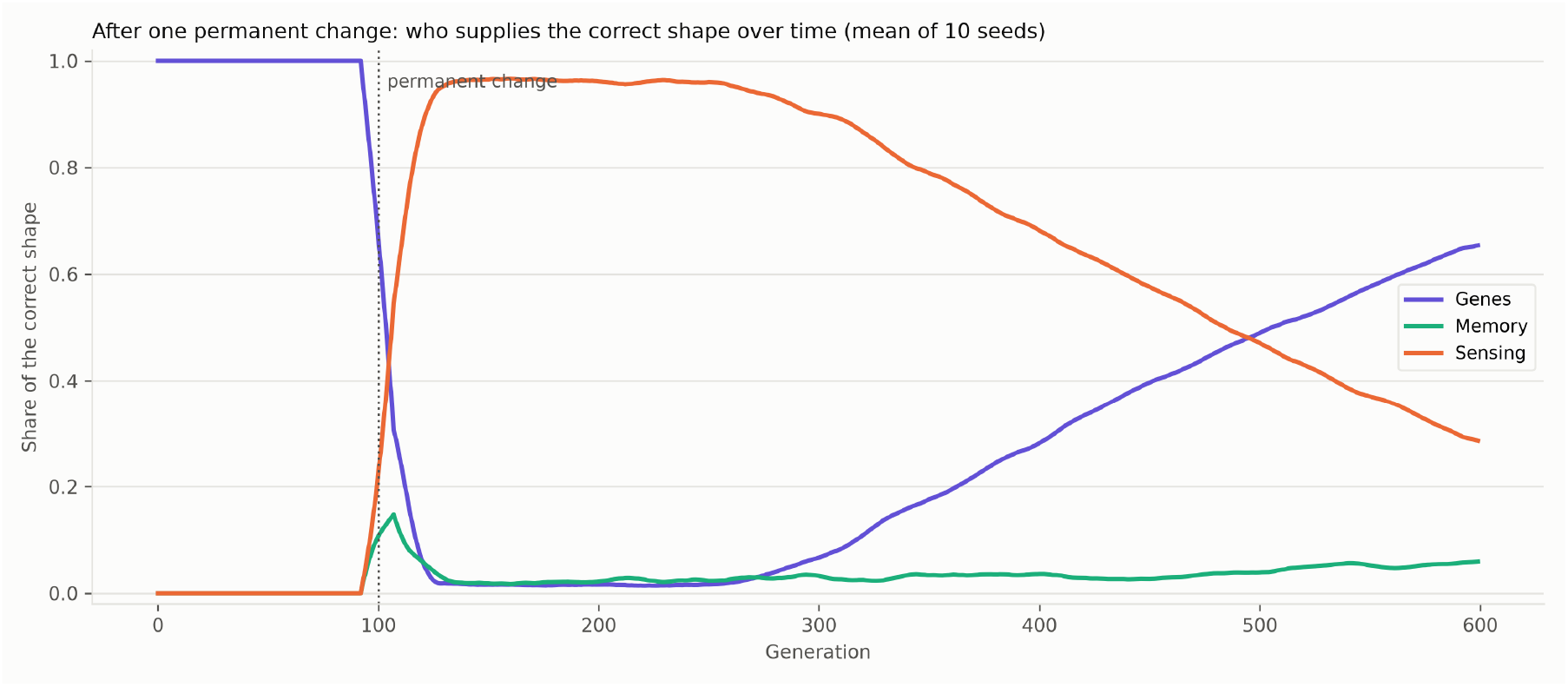
Version 16. Share of the correct form supplied by genes, memory and sensing over time (mean of 10 seeds). After the permanent change at generation 100, sensing takes over almost at once; genes overtake it through selection roughly 340 to 450 generations later.

### 4.5 A fresh pre-specified re-test of the handover (version 22)

The version 16 model was rerun unchanged on 20 new seeds (1000 to 1019), with the crossover rule, the late window (generations 550 to 599) and the thresholds fixed in writing beforehand. The handover was **replicated** (Figure 2). Genes overtook sensing in 20 of 20 seeds, at generations 452 to 588 (median 496, IQR 481 to 532; p = 1.9×10^−6^); genes were ahead of sensing in the late window in 19 of 20 seeds (p = 4.0×10^−5^; mean shares: genes 0.60, memory 0.05, sensing 0.35); and sensing supplied a mean of 0.94 of the correct form at generation 125. The dependence on costs and inheritance was only partly reproduced: genes led with halved costs (8 of 10 seeds) and with 20% carry-over (10 of 10), and sensing led with no carry-over (8 of 10), but with doubled costs the lead was split (5 of 10 each), where version 16 had given sensing 8 of 10. The dependence of the handover on cost is therefore weaker than version 16 suggested.

**Figure 2.**
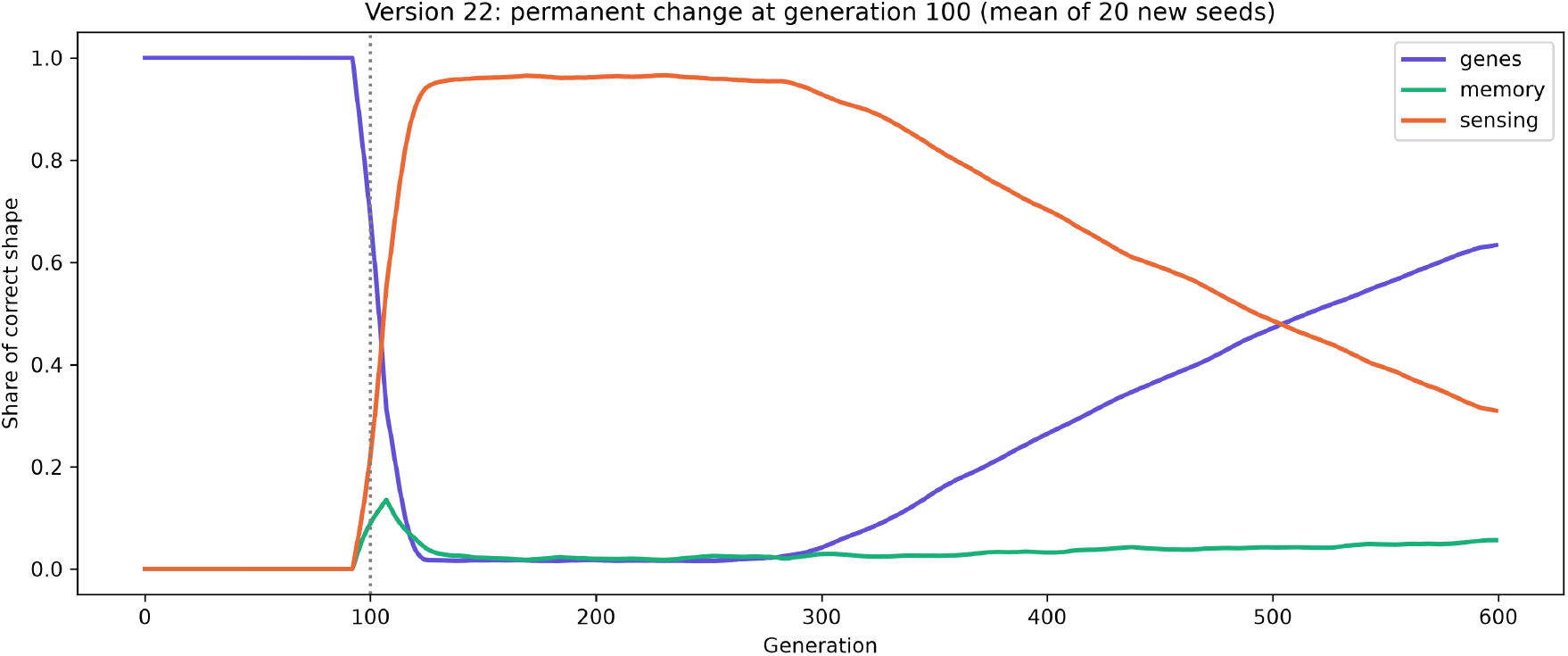
Version 22. Share of the correct form supplied by genes, memory and sensing after the permanent change at generation 100 (mean of 20 new seeds).

## 5. Results, Part 2: energy-switched release of concealed variation (versions 17–21)

The release mechanism behaved as predicted in every version, but its net value reversed between versions 19 and 20. Version 21 (section 5.5) shows that the reversal came from back mutation, which caps the harmful load, and not from population size.

| **Version** | **Population** | **Release vs no control (mean fitness)** | **Seeds favouring release** | **Sign-test p** | **Pre-specified verdict** |
|---|---|---|---|---|---|
| 18 | 500 | 0.603 vs 0.629 | 1 of 20 | 4.0×10<sup>−5</sup> | Net cost |
| 19 | 500 | 0.103 vs 0.122 | 3 of 20 | 0.003 | Net cost (12 of 13 robustness settings) |
| 20 | 2,000 | 0.632 vs 0.489 | 20 of 20 | 1.9×10<sup>−6</sup> | Net benefit (8 of 8 robustness settings) |

### 5.1 Concealment and release (versions 17–18)

With neutral shape mutations (version 17), controlled populations held 3.3 times the genetic variance of uncontrolled ones at the moment of environmental change (0.55 against 0.16; 20 of 20 seeds). An energy shortfall raised visible variation fourteen-fold, and released populations reached half-way to the new optimum in a median of 4 generations against 6 without control, while control that stayed strong took 43 generations. When 80% of mutations were harmful (version 18), control concealed 5.9 times the harmful load (2.25 against 0.38; 20 of 20), and release cut the defect component of fitness to 0.18 against 0.69. Release still sped adaptation (6 against 10 generations) but lost overall (0.603 against 0.629; lower in 19 of 20 seeds), with robustness on a knife edge (release won in 5 of 10 settings).

### 5.2 Duration and re-locking (version 18)

Mean fitness after the change was 0.573 for a 5-generation shortfall, 0.603 for 20 generations, 0.563 for 50 and 0.274 for a shortfall that never ended. Release followed by re-locking around the new form outperformed release with control fixed on the old form (0.603 against 0.445; 20 of 20). Release at random times was worse than perfectly timed release in all 20 seeds and worse than no control (Figure 3).

**Figure 3.**
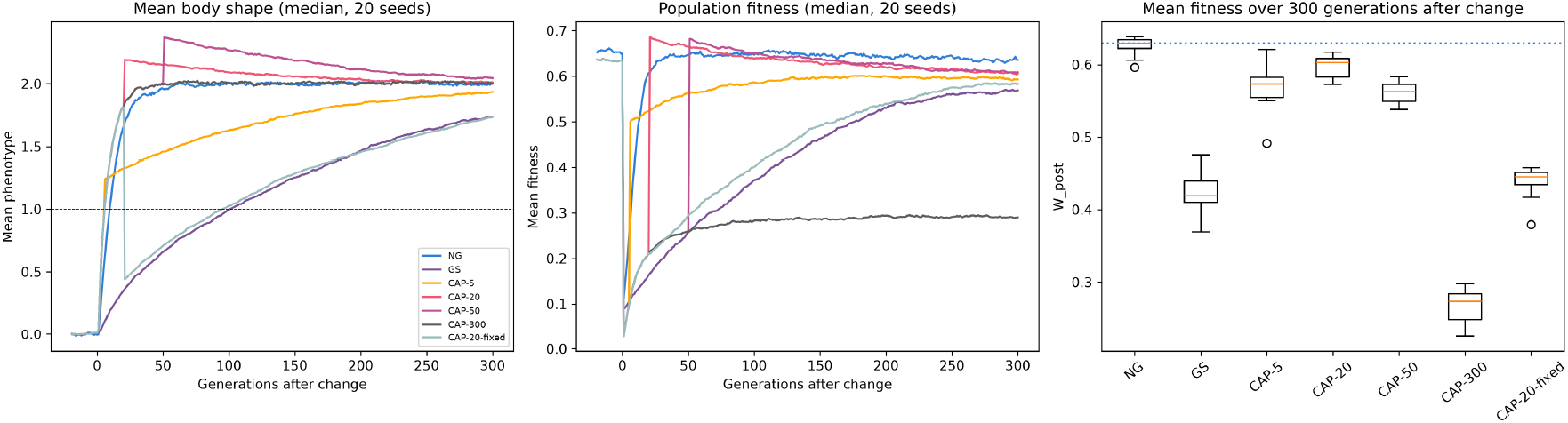
Version 18. Left: mean form after the change. Centre: population fitness. Right: mean fitness over 300 generations; release adapts fastest but stays below no control, and a permanent shortfall is worst. NG, no control; GS, control never stressed; CAP-n, control with a shortfall of n generations at the change (CAP-300 is effectively permanent); CAP-20-fixed, a 20-generation shortfall after which control stays locked on the old form.

### 5.3 Published mutation figures, population 500 (version 19)

With published mutation parameters rescaled to a population of 500 and ten successive environmental changes, control concealed about five times the harmful load (9.7 against 2.0; 20 of 20). Release sped catch-up after each change (7.6 against 11.3 generations; 20 of 20) but was a net cost even when perfectly timed (0.103 against 0.122; lower in 17 of 20), and randomly timed release was costlier still (0.091; 20 of 20). During each release, mean fitness fell close to zero for about 20 generations (Figure 4).

**Figure 4.**
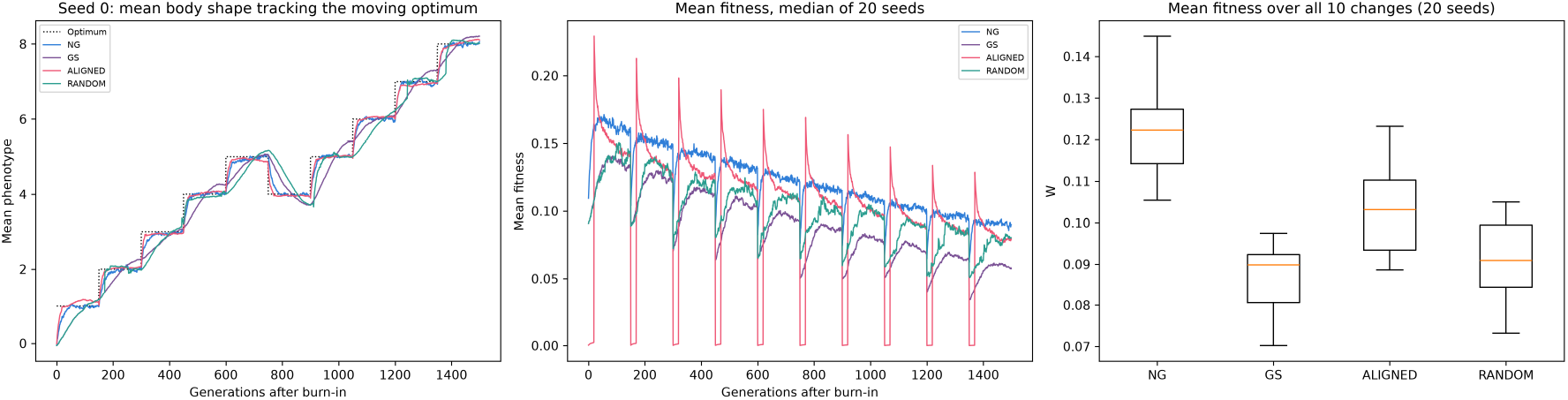
Version 19. Left: one seed tracking the moving optimum. Centre: median fitness over 20 seeds, collapsing during each release. Right: mean fitness across ten changes. NG, no control; GS, control never stressed; ALIGNED, a 20-generation shortfall at each change; RANDOM, shortfalls at random times unrelated to the changes.

### 5.4 Larger populations with back mutation (version 20)

Under the conditions identified by Masel (2005, 2006), the result reversed: with 2,000 individuals and back mutation, perfectly timed release outperformed no control in all 20 seeds (0.632 against 0.489; median paired difference +0.143, IQR +0.142 to +0.144; sign test p = 1.9×10^−6^) and in all 8 robustness settings. Post-hoc diagnostics on 10 seeds showed that control without release already outperformed no control (0.571 against 0.489; 10 of 10), and that release added a further gain (0.632 against 0.571; 10 of 10); lower developmental noise alone did not reproduce the gain (0.373). Fitness still fell during each release (mean 0.33, minimum 0.10) but recovered to 0.68 between releases. In a two-trait valley requiring coordinated change, no group crossed in 1,000 generations, and with a shallower valley uncontrolled populations crossed more often (9 of 10 against 4 of 10 seeds), so the predicted multi-step advantage of release was not reproduced (Figure 5).

**Figure 5.**
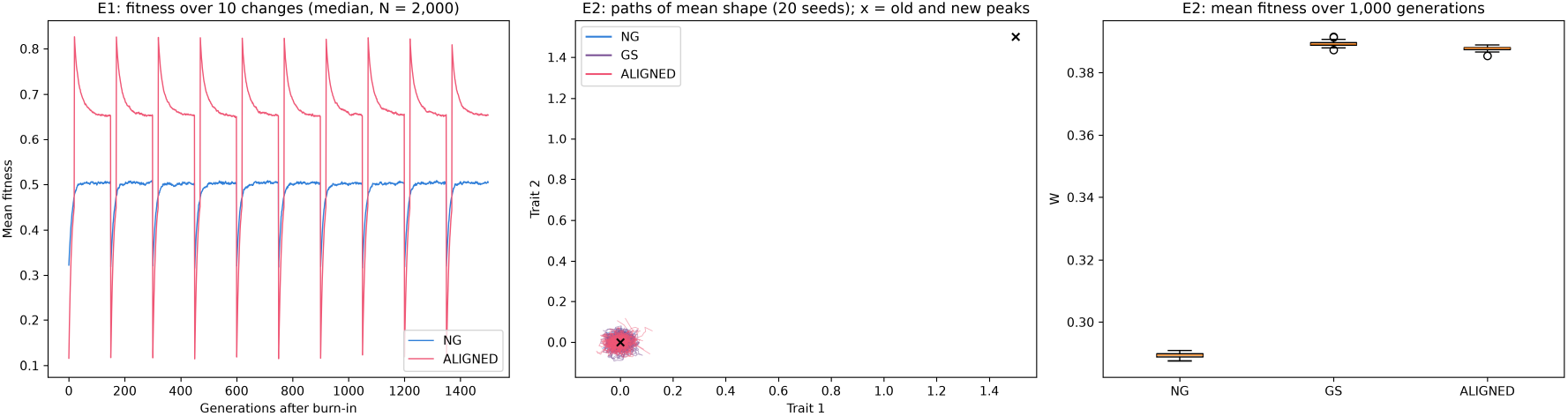
Version 20. Left: fitness over ten changes; release stays above no control. Centre: paths of mean form in the two-trait valley; no group reaches the new peak. Right: mean fitness in the valley experiment. Groups as in Figure 4 (NG, GS, ALIGNED).

### 5.5 Separating population size from rescaling and back mutation (version 21)

Between versions 19 and 20 three things changed together: population size, the rescaling factor Q, and back mutation. Version 21 reused the version 20 model unchanged and crossed them (20 seeds per cell; ten shifts of the optimum; groups: no control, control never stressed, control with an aligned shortfall). It first reproduced version 20 exactly (per-seed values matched to 10^−16^) and reproduced the version 19 cost as a statistical check (release below no control in 20 of 20 seeds at N = 500, Q = 20, no back mutation).

The pre-specified verdict was **Outcome 2: population size is not sufficient** (robustness 6 of 6). With Q = 5 and back mutation at 10% of the forward rate, release was a net benefit at every population size from 250 to 4,000 (release below no control in 0 of 20 seeds in each). At N = 500, removing back mutation alone gave a net cost (16 of 20 seeds; sign test p = 0.012, uncorrected; median difference −0.007), whereas changing Q alone did not (0 of 20).

| **Cell (N, Q, back mutation)** | **Seeds with release below no control** | **Sign-test p** | **Median difference, release minus no control (IQR)** | **Class** | **Median fitness: no control / control only / release** |
|---|---|---|---|---|---|
| 250, 5, 0.1 | 0 | 1.9×10<sup>−6</sup> | +0.079 (+0.060 to +0.085) | benefit | 0.469 / 0.473 / 0.548 |
| 500, 5, 0.1 | 0 | 1.9×10<sup>−6</sup> | +0.112 (+0.109 to +0.116) | benefit | 0.481 / 0.518 / 0.592 |
| 2,000, 5, 0.1 | 0 | 1.9×10<sup>−6</sup> | +0.143 (+0.142 to +0.144) | benefit | 0.489 / 0.570 / 0.632 |
| 4,000, 5, 0.1 | 0 | 1.9×10<sup>−6</sup> | +0.145 (+0.144 to +0.145) | benefit | 0.489 / 0.585 / 0.634 |
| 500, 20, 0.1 | 0 | 1.9×10<sup>−6</sup> | +0.111 (+0.110 to +0.112) | benefit | 0.412 / 0.483 / 0.523 |
| 500, 5, none | 16 | 0.012 | −0.007 (−0.013 to −0.002) | cost | 0.091 / 0.066 / 0.085 |
| 500, 20, none | 20 | 1.9×10<sup>−6</sup> | −0.019 (−0.023 to −0.009) | cost | 0.123 / 0.088 / 0.105 |
| 1,000, 20, none | 15 | 0.041 | −0.009 (−0.018 to −0.002) | mixed | 0.184 / 0.152 / 0.174 |
| 2,000, 5, none | 11 | 0.824 | −0.001 (−0.008 to +0.007) | mixed | 0.218 / 0.186 / 0.217 |
| 2,000, 20, none | 12 | 0.503 | −0.003 (−0.007 to +0.004) | mixed | 0.234 / 0.214 / 0.232 |
| 4,000, 20, none | 1 | 4.0×10<sup>−5</sup> | +0.016 (+0.009 to +0.020) | benefit | 0.265 / 0.259 / 0.281 |

(Medians over 20 seeds; the sign test cannot give a p-value below 1.9×10^−6^ with 20 seeds. The remaining cells, including N = 1,000 and 2,000 at Q = 20 with back mutation, are in the repository files `v21_verdicts.json` and `v21_per_seed.json`; Figure 6 shows every cell.)

**Figure 6.**
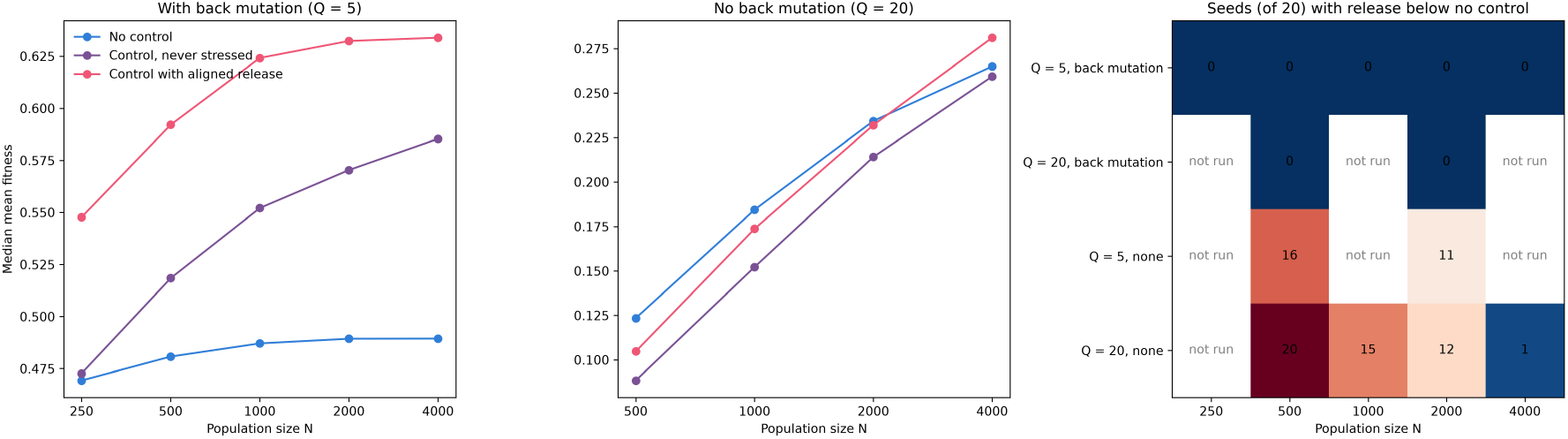
Version 21. Left: median fitness by population size with back mutation (Q = 5). Centre: the same without back mutation (Q = 20). Right: number of seeds (of 20) in which release is below no control, in every cell run; blue cells are net benefit, red cells net cost, and “not run” cells were not part of the design.

The sizes of the effects differ greatly. Without reversal the cost was small (median difference −0.007 and −0.019 fitness at N = 500), and the p-values of the cost cell and the nearest mixed cell are modest and uncorrected (0.012 at N = 500, Q = 5; 0.041 at N = 1,000, Q = 20), so they are weak evidence of a cost on their own; with reversal the benefit was large (+0.079 to +0.145). The classes in the table rest on the pre-specified seed-count thresholds, not on these p-values.

Three further observations follow, the first two from measures fixed in advance but outside the settle rule, the third not pre-specified. First, release was never below control left unstressed, in any of the 13 cells: the net cost seen without back mutation lies in the control itself (control alone was below no control in 20 of 20 seeds at N = 500, 1,000 and 2,000), and release recovers part of that cost. Second, with back mutation, control alone already beat no control from N = 1,000 upwards. Third, without back mutation a population-size gradient remains (at Q = 20, release was below no control in 20, 15, 12 and 1 of 20 seeds at N = 500, 1,000, 2,000 and 4,000), so size matters when harmful load cannot be reversed, but it is not what reverses the result when it can. In this model back mutation caps the harmful load; real back mutation per site is far rarer, so this factor may stand in for another process that lets accumulated harmful load shrink, which version 21 does not test.

## 6. Discussion

### 6.1 One picture from two parts

The two parts of the programme describe a single system seen at two moments. In ordinary times, energy-funded control holds the body to a working form, absorbs change faster than genes can, and is trimmed back wherever it is not earning its cost. Because it corrects the body regardless of genotype, it conceals genetic variation from selection, and selection meanwhile writes lasting changes into the genes over hundreds of generations (Part 1).

When energy runs short, the same control weakens without any decision being made, and the concealed variation is exposed all at once (Part 2). The emphasis of the original idea therefore shifts from “steering guides evolution” to “control buys time while selection rewrites the genes, and its failure under energy shortage is an evolutionary gamble”.

### 6.2 What decides the gamble

An initial interpretation was that the reversal between versions 19 and 20 was a population-size effect through the drift barrier: selection acts on a mutation only when its effect exceeds roughly the reciprocal of population size, so control hides many harmful mutations below that threshold in small populations and a cleaner store survives in large ones (Masel, 2005, 2006). Version 21 tested that explanation directly and did not support it as the main cause. Changing population size from 250 to 4,000 left release a net benefit as long as harmful mutations could revert, and removing reversion reversed the result at N = 500 regardless of the rescaling factor. What the model shows is that the sign depends first on whether the hidden harmful load is kept in check, and only second, and without reversion, on population size. The measured hidden-load ratio (control against no control) was about 3 with reversion and 4 to 5 without, at all sizes tested. The drift-barrier account remains possible for the residual size gradient seen without reversion, but it is not demonstrated here, and a model with a different way of limiting harmful load might give a different dependence on size.

### 6.3 Relation to existing work

Much of the Part 1 behaviour is anticipated by plasticity theory and by earlier models of the Baldwin effect (Lande, 2009): the dependence of strategy on timescale and cue reliability (Botero et al., 2015), plasticity first and genetic assimilation later (Scheiner et al., 2017), and the integration of several information sources (McNamara et al., 2016). Concealment and stress-triggered release are established for cryptic genetic variation and Hsp90 (Paaby and Rockman, 2014; Rutherford, Hirate and Swalla, 2007; Sgrò et al., 2010), and occur in nature (Rohner et al., 2013). The pattern of harmful load in small populations agrees with genomic work on island foxes under an additive model, although strongly deleterious recessive alleles were purged there (Robinson et al., 2018), a case our model does not represent.

The closest published neighbours to the central mechanism are these. Shreesha and Levin (2023) and Hartl, Risi and Levin (2024) show in simulation that cellular competency during development masks the raw fitness of genomes from selection, but their competency carries no energy cost in the published model, and neither treats release of concealed variation. Rutherford et al. (2007) propose that stress overwhelms the buffering capacity of the chaperone Hsp90 so that cryptic variation is expressed, which is a capacity-limit mechanism close in spirit to ours; Chen and Wagner (2012) show that reduced Hsp90 expression lowers fecundity, longevity and the buffering of cryptic deleterious variation in wild flies. Le Rouzic, Álvarez-Castro and Hansen (2013) argue that stress-linked decanalization may be an automatic side effect rather than an adaptation, as in our model. Nelson and Masel (2018) show that purging of deleterious cryptic sequences can arise without a dedicated capacitor, which is mechanistically near our back-mutation result. Trotter et al. (2014) show that capacitors can make crossing wide fitness valleys common; our two-trait valley test did not reproduce this and is inconclusive.

Following Masel and Siegal (2009), we call our mechanism a phenotypic capacitor rather than an evolutionary capacitor, because the model shows release of concealed variation but does not show that the released variation is later assimilated.

### 6.4 What may be new

Three elements were not found in the sources we examined: (i) a single model in which bioelectric control, memory, sensing and genes share one energy budget, with control trimmed where it does not pay; (ii) a developmental control system, interpreted here as a bioelectric network, acting as a phenotypic capacitor whose strength is set by an energy budget, so that energy shortfall is the switch; and (iii) a model result that the value of such an energy-switched capacitor depends mainly on whether the harmful load it conceals is kept in check, with the population-size dependence of capacitance theory appearing only weakly and only where harmful mutations cannot revert. A search of Europe PMC (which indexes PubMed, preprints and other records) on 20 October 2026, with seven Boolean queries on these elements and forward-citation checks of Masel (2005), Shreesha and Levin (2023) and Hartl et al. (2024), found no record that sets the strength of a capacitor by an energy budget or treats a bioelectric network as a capacitor (Supplementary Note 1). The search did not cover Web of Science, Scopus or Google Scholar and cannot certify absence. The contribution is therefore best described as a specific, model-based variant of established ideas.

### 6.5 Limitations

- Versions 15 and 16 were recovered from the earlier session and version 16 reproduced exactly; the code for earlier Part 1 versions (1 to 14) has not yet been recovered, so their results rest on the original runs. The central handover result was identified after scoring and was then replicated on new seeds in version 22 (section 4.5), under the same model and environment pair.
- Several results from exploratory voice conversations with the AI assistant were never reproduced in text and are excluded.
- The models are minimal: few traits, additive effects, fixed control strength, and stress that affects the whole population at once.
- Version 21 separated the three parameters that changed between versions 19 and 20 and found that back mutation, not population size, reversed the result; but back mutation here is a simple cap on harmful load, population sizes stop at 4,000, and the two-trait valley experiment was not repeated.
- Mutation figures come from different species, and one parameter (the mean harmful effect) was fitted to a published summary.
- The behaviour route, in which behaviour changes the environment and genes follow, worked in only 5 of 10 runs and depended on a chosen bonus value.
- The models contain no voltages, ion channels or tissue; control strength is an abstract parameter, and its identification with a bioelectric network is an interpretation, not something the simulations test.
- No part of the bioelectric mechanism has been measured in a living organism.

### 6.6 Testable predictions

In developing embryos, such as zebrafish or *Xenopus*, from several genetic lines, with membrane voltage measured by fluorescent reporters (Adams and Levin, 2012):

1. Mild energy restriction lowers membrane voltage and widens the spread of body forms.
2. Differences between genetic lines widen under restriction, the signature of released concealed variation rather than random noise.
3. Most newly exposed forms are defects, and long restriction is worse than short.
4. Holding membrane voltage steady pharmacologically during restriction cancels these effects.

At the population level, the net fitness effect of repeated release should depend mainly on how much harmful load a population can shed, and on population size chiefly where harmful load cannot be reversed; and after a permanent environmental change, genetic adaptation should follow physiological adaptation on a timescale of hundreds of generations.

## 7. Conclusion

Twenty-two versions of pre-specified simulation do not support the claim that developmental control steers evolution, but they support a more specific account of what energy-funded control does. It absorbs change quickly while selection writes lasting change into the genes over hundreds of generations, it is kept only where it pays, and it conceals rather than directs genetic variation. When energy runs short it releases what it concealed, an evolutionary gamble whose outcome depends on the duration of the shortfall, on whether control re-locks around the new form and, above all, whether the concealed harmful load is kept in check, with population size mattering more weakly. Measuring the mechanism in developing embryos, testing other ways of limiting harmful load, repeating the multi-step case and completing a systematic literature search are the necessary next steps.

## Supporting information

Supplementary Note 1: literature search

Supplementary Table 1: spread and sign-test statistics

## Declarations

### Code and data availability

Simulation code, pre-specifications (named pre-registrations in the repository) with change logs, verdict files, figures and audit records are deposited at https://github.com/excoms/steering-before-selection and archived on Zenodo (version 1.0.0, doi:10.5281/zenodo.23920330; all versions, doi:10.5281/zenodo.23920329). Every analysis uses fixed random seeds. A full rerun of the analyses for versions 16 to 20 on 20 October 2026 (Python 3.13.2, NumPy 2.5.4) reproduced the shipped result files exactly or to floating-point precision, and version 21 reproduced the per-seed values of version 20 exactly as a built-in check. **Licences**. The code is released under the MIT licence; the text, figures and result files under CC BY 4.0. The preprint is posted under CC BY.

### Supplementary material

Supplementary Note 1 records the literature search (queries, hit counts, citation checks and limits). Supplementary Table 1 gives spread and sign-test statistics for every comparison.

### Pre-specification

From version 11 onwards, predictions, thresholds and failure conditions were written before each version was run; versions 1 to 10 were exploratory. Departures are recorded in each version’s change log and identified as post-hoc in the text. The pre-specifications were written in the project files before each run, with the code and rule files fixed by hash for versions 21 and 22, but they were not lodged with an external registry, so their timing rests on the authors’ records and file history and not on independent time-stamping.

Results obtained only in exploratory voice conversations with the AI assistant and never reproduced are excluded.

### Use of AI assistance

The simulation code for versions 1 to 20, the analyses and a first draft of this manuscript were produced with Claude (Anthropic) in claude.ai chat sessions, using Claude Opus 5.5, working under the authors’ direction. Versions 21 and 22, the statistical analysis, the checks of numbers and references, and the revision of the manuscript were produced with Claude Sonnet 5.5 (claude-sonnet-5-5) in Claude Code. The authors developed the idea together and reviewed and approved the conclusions; M.Q. directed the work. The authors take responsibility for all content.

### Author contributions

The authors developed the idea together. M.Q. directed the development of the work; F.F.Q. supported it.

### Review status

This preprint was posted without review by anyone outside the author team. Comments received after posting will be addressed in revised versions under the same DOI.

