## Supplementary Note 1: literature search for "Steering Before Selection: Energy-Limited Developmental Control as a Phenotypic Capacitor, a Pre-Specified Simulation Study"

Run on 20 October 2026 on Europe PMC (https://europepmc.org), which indexes PubMed, preprint servers (including bioRxiv) and other life-science records, through its public REST search interface. The aim was to look for prior work on the three elements the paper treats as possibly new: (i) a model with control, memory, sensing and genes sharing one energy budget; (ii) a capacitor whose strength is set by an energy budget, so that energy shortfall is the switch, and a bioelectric network as the capacitor; (iii) a net value of release that depends mainly on whether harmful load is kept in check.

### Queries and results

| # | Query (Boolean, as run) | Records | Screened | Relevant to the three elements |
| --- | --- | --- | --- | --- |
| 1 | (“evolutionary capacitor” OR “evolutionary capacitance” OR “phenotypic capacitor” OR “phenotypic capacitance” OR “Hsp90 capacitor”) AND (energy OR ATP OR metabolic OR “energy budget” OR “energy cost” OR starvation) | 98 | all 98 | None. Hits are Hsp90 and capacitor background. |
| 2 | (“cryptic genetic variation” OR the capacitor terms of query 1) AND (bioelectric* OR “membrane potential” OR “membrane voltage” OR “resting potential”) | 4 | all 4 | None (plant systems biology, signalling reviews). |
| 3 | (“evolutionary capacitance” OR “cryptic genetic variation” OR “evolutionary capacitor”) AND (“back mutation” OR reversion OR “reverse mutation”) AND (“population size” OR “effective population size”) | 14 | all 14 | Masel 2005 and 2006, the [PSI] capacitor papers, Nature Communications 2025 cavefish study. None treats reversal of harmful load as the factor that sets the sign of release. |
| 4 | bioelectric* AND (evolution OR evolvability OR “natural selection”) AND (simulation OR “computational model” OR “in silico”) | 1,110 | first 100 by relevance | Levin-group simulations: Hartl et al. 2024; Pio-Lopez, Hartl and Levin 2025 (aging as loss of goal-directedness). None has an energy-switched capacitor. |
| 5 | (“morphogenetic competency” OR “cellular competency” OR “developmental competency”) AND (evolution OR evolvability OR selection) | 256 | first 100 by relevance | Shreesha and Levin 2023; Hartl et al. 2024; Levin 2023 (Darwin’s agential materials); Pio-Lopez et al. 2023 (scaling of goals). No energy cost in the published models. |
| 6 | (canalization OR canalisation OR “phenotypic buffering” OR “developmental buffering” OR robustness) AND (“energetic cost” OR “energy cost” OR “metabolic cost” OR “cost of buffering” OR “cost of robustness” OR “cost of canalization”) AND (evolution) AND (model OR simulation) | 2,187 | first 100 by relevance | None (results dominated by unrelated engineering papers). |
| 7 | (decanalization OR decanalisation OR “release of cryptic” OR “cryptic variation”) AND (stress) AND (energy OR nutritional OR starvation OR “resource limitation”) | 139 | first 100 by relevance | Competition and cryptic variation in tadpoles (2026); a bacterial Rho capacitor (2025). Neither is an energy-limited capacitor model. |

### Forward-citation checks (Europe PMC “cited by”)

| Paper | Citing records indexed | Titles scanned for energy, bioelectric, capacitor, competency, cryptic, buffering, canalization, cost |
| --- | --- | --- |
| Masel 2005, *Genetics* | 50 | 22 matched; none proposes an energy-switched capacitor or a bioelectric capacitor. Nearest: Trotter et al. 2014 (valley crossing), Nelson and Masel 2018 (emergent capacitance), reviews of robustness and evolvability. |
| Shreesha and Levin 2023, *Entropy* | 16 | 2 matched (a bioelectricity-and-aging review; Levin 2023 review). None adds an energy cost or a capacitor. |
| Hartl, Risi and Levin 2024, *Entropy* | 9 | 0 matched. |

### Related work found and used in the paper

Shreesha and Levin 2023; Hartl et al. 2024; Rutherford et al. 2007; Chen and Wagner 2012; Le Rouzic et al. 2013; Nelson and Masel 2018; Trotter et al. 2014; Masel and Siegal 2009; Sgrò et al. 2010; Masel 2005 and 2006.

### Limits

- One database. Web of Science, Scopus and Google Scholar were not searched; Europe PMC citation counts are lower than theirs.
- For queries 4 to 7 only the first 100 records by relevance were screened, by title (and, for six candidates, abstract). Absence from those lists does not prove absence from the literature.
- Keyword queries miss work that uses different terms (for example “robustness switch”, “buffering capacity”, “resource-dependent canalization”).
- A search that finds nothing is weak evidence of absence. The paper therefore describes its contribution as a specific, model-based variant of established ideas and does not claim priority.
