## Supplementary Table 1: spread and sign-test statistics for "Steering Before Selection: Energy-Limited Developmental Control as a Phenotypic Capacitor, a Pre-Specified Simulation Study"

### Supplementary Table 1. Spread and sign tests for the main comparisons

Medians of per-seed paired differences with interquartile ranges (IQR), and exact two-sided sign-test p-values (null probability 0.5) for the number of seeds in which one group is above the other. These are descriptive: the verdicts in the paper were fixed in advance by seed-count thresholds, seeds are simulation replicates, and no correction is made for multiple comparisons. Per-seed values were saved for versions 20 and 21 only; for versions 16 to 19 the table gives seed counts.

#### Version 21, all 13 cells (20 seeds each; W is mean fitness over the ten changes, excluding the energy cost)

| Cell | Release below no control | p | Median difference release − no control (IQR) | Release below control only | p | Control only below no control | p |
| --- | --- | --- | --- | --- | --- | --- | --- |
| N1000_Q20_b0.0 | 15/20 | 0.041 | −0.009 (−0.018 to −0.002) | 0/20 | 1.9×10⁻⁶ | 20/20 | 1.9×10⁻⁶ |
| N1000_Q5_b0.1 | 0/20 | 1.9×10⁻⁶ | +0.136 (+0.132 to +0.139) | 0/20 | 1.9×10⁻⁶ | 0/20 | 1.9×10⁻⁶ |
| N2000_Q20_b0.0 | 12/20 | 0.503 | −0.003 (−0.007 to +0.004) | 0/20 | 1.9×10⁻⁶ | 20/20 | 1.9×10⁻⁶ |
| N2000_Q20_b0.1 | 0/20 | 1.9×10⁻⁶ | +0.113 (+0.112 to +0.113) | 0/20 | 1.9×10⁻⁶ | 0/20 | 1.9×10⁻⁶ |
| N2000_Q5_b0.0 | 11/20 | 0.824 | −0.001 (−0.008 to +0.007) | 0/20 | 1.9×10⁻⁶ | 20/20 | 1.9×10⁻⁶ |
| N2000_Q5_b0.1 | 0/20 | 1.9×10⁻⁶ | +0.143 (+0.142 to +0.144) | 0/20 | 1.9×10⁻⁶ | 0/20 | 1.9×10⁻⁶ |
| N250_Q5_b0.1 | 0/20 | 1.9×10⁻⁶ | +0.079 (+0.060 to +0.085) | 0/20 | 1.9×10⁻⁶ | 9/20 | 0.824 |
| N4000_Q20_b0.0 | 1/20 | 4.0×10⁻⁵ | +0.016 (+0.009 to +0.020) | 0/20 | 1.9×10⁻⁶ | 17/20 | 0.003 |
| N4000_Q5_b0.1 | 0/20 | 1.9×10⁻⁶ | +0.145 (+0.144 to +0.145) | 0/20 | 1.9×10⁻⁶ | 0/20 | 1.9×10⁻⁶ |
| N500_Q20_b0.0 | 20/20 | 1.9×10⁻⁶ | −0.019 (−0.023 to −0.009) | 0/20 | 1.9×10⁻⁶ | 20/20 | 1.9×10⁻⁶ |
| N500_Q20_b0.1 | 0/20 | 1.9×10⁻⁶ | +0.111 (+0.110 to +0.112) | 0/20 | 1.9×10⁻⁶ | 0/20 | 1.9×10⁻⁶ |
| N500_Q5_b0.0 | 16/20 | 0.012 | −0.007 (−0.013 to −0.002) | 0/20 | 1.9×10⁻⁶ | 20/20 | 1.9×10⁻⁶ |
| N500_Q5_b0.1 | 0/20 | 1.9×10⁻⁶ | +0.112 (+0.109 to +0.116) | 0/20 | 1.9×10⁻⁶ | 1/20 | 4.0×10⁻⁵ |

#### Version 20

| Experiment | Release below no control | p | Median difference (IQR) |
| --- | --- | --- | --- |
| E1, repeated changes | 0/20 | 1.9×10⁻⁶ | +0.143 (+0.142 to +0.144) |
| E2, valley experiment | 0/20 | 1.9×10⁻⁶ | +0.098 (+0.097 to +0.099) |

#### Versions 16 and 22: generation at which genes overtake sensing

| Version | Seeds crossing | p | Median (IQR) | Range |
| --- | --- | --- | --- | --- |
| 16 | 10/10 | 0.002 | 489 (462 to 510) | 438 to 549 |
| 22 | 20/20 | 1.9×10⁻⁶ | 496 (481 to 532) | 452 to 588 |

Version 22, genes ahead of sensing in the late window (generations 550 to 599): 19/20 seeds, p = 4.0×10⁻⁵.

#### Versions 17 to 19: seed counts only

| Version | Claim | Seeds | p |
| --- | --- | --- | --- |
| v17 | guard holds more genetic variance than no guard | 20/20 | 1.9×10⁻⁶ |
| v17 | shortfall raises visible variation | 20/20 | 1.9×10⁻⁶ |
| v17 | released populations not slower to half-way | 20/20 | 1.9×10⁻⁶ |
| v18 | control conceals at least 3 times the harmful load | 20/20 | 1.9×10⁻⁶ |
| v18 | release lowers the defect component of fitness | 20/20 | 1.9×10⁻⁶ |
| v18 | release below no control (net cost) | 19/20 | 4.0×10⁻⁵ |
| v18 | random timing below perfectly timed release | 20/20 | 1.9×10⁻⁶ |
| v18 | re-lock above fixed control | 20/20 | 1.9×10⁻⁶ |
| v19 | guard hides harmful load | 20/20 | 1.9×10⁻⁶ |
| v19 | release faster catch-up | 20/20 | 1.9×10⁻⁶ |
| v19 | release below no control (net cost, perfectly timed) | 17/20 | 0.003 |
| v19 | random timing below no control | 20/20 | 1.9×10⁻⁶ |
